# Blunted fear-evoked striatal dopamine release in individuals with a family history of psychosis

**DOI:** 10.64898/2026.08.24.746811

**Authors:** Rami Hamati, Cecelia Shvetz, Bianca Chidiac, Hussein Bdair, Katie Dinelle, Daphne J. Holt, Clifford M. Cassidy, Lauri Tuominen

## Abstract

While excess tonic dopamine signalling is a hallmark of schizophrenia and psychotic disorders, it has been difficult to reconcile with dopamine-dependent learning deficits seen in schizophrenia. Excess spontaneous activity of tonic dopamine neurons, coupled with reduced coordinated activity of phasic dopamine neurons, may explain the observed discrepancy between increased tonic signalling and impaired learning. Although intriguing, this chaotic dopamine hypothesis lacks empirical support. In the current study, Pavlovian fear conditioning is used to test this hypothesis in healthy individuals with and without a family history of psychosis using simultaneous [^11^C]raclopride PET/fMRI. In 16 healthy individuals without a family history of psychosis, we first show that fear conditioning releases dopamine and link this release to BOLD responses. We then report that in 12 first-degree relatives of individuals with psychotic disorders, this adaptive dopamine release in the posterior caudate is lacking, despite no differences in behavioural learning. Furthermore, reduced dopamine release is associated with increased self-reported paranoid thinking, but not with anhedonia. These findings provide novel *in vivo* evidence supporting the chaotic dopamine hypothesis, suggesting that an adaptive, stimulus-driven dopamine release is lacking in psychotic disorders and may contribute to positive symptoms like paranoia.

## Introduction

Schizophrenia is a disorder of excess tonic dopamine signaling [1]–[3]. Positron emission tomography (PET) studies have demonstrated elevated dopamine synthesis capacity, higher dopamine release in response to amphetamines, and increased synaptic dopamine levels in the striatum in individuals with schizophrenia [4]. Importantly, excess dopamine signaling extends to individuals with other psychotic disorders, clinical risk for psychosis, first degree relatives, and those with higher polygenic risk score for psychosis [5]–[8].

Yet, excess tonic dopamine signaling has been difficult to reconcile with findings of impaired dopamine-dependent learning in psychotic disorders. For example, individuals with psychotic disorders show reduced behavioural responses to reward predicting cues [9], [10], which implies that individuals with psychotic disorders have reduced dopamine signalling during the task. In functional magnetic resonance imaging (fMRI) studies using reward tasks, individuals with psychotic disorders have blunted striatal BOLD responses during anticipation of reward, and to reward prediction errors [11]–[14], which again implies reduced dopamine signalling during the tasks. In addition to reward learning, individuals with schizophrenia exhibit impaired discriminative fear conditioning, another dopamine dependent process [15]–[17]. Thus, if schizophrenia was simply a disorder of appropriately synchronized but excessive dopamine signaling, then the performance in dopamine dependent tasks should be better, not worse.

The chaotic dopamine hypothesis has emerged as a prominent theory trying to resolve this discrepancy [18], [19]. It postulates an increase in spontaneous dopamine release that contributes to higher tonic levels of dopamine and a reduction of adaptive, stimulus-driven phasic dopamine release in schizophrenia. Furthermore, the hypothesis predicts that increased spontaneous dopamine release is associated with positive symptoms, while decreased adaptive dopamine release is associated with negative symptoms. While compelling, so far there has been limited evidence supporting the chaotic dopamine hypothesis.

In this study, we use Pavlovian fear conditioning to directly test the chaotic dopamine hypothesis. Pavlovian fear conditioning requires adaptive, stimulus-driven dopamine release to form associations between aversive and neutral stimuli in rodents [16]. However, definitive evidence that fear conditioning releases dopamine in humans is lacking [16]. Therefore, we first seek to demonstrate that fear conditioning leads to detectable dopamine release in the striatum in individuals without family history of psychosis and explore the link between dopamine release and other fear responses. We measure dopamine release, neural, physiological, and behavioural responses simultaneously with [^11^C]raclopride PET/fMRI, skin conductance, and behavioural ratings. Then, we set out to test the chaotic dopamine hypothesis in a group of first-degree relatives (FDRs) of individuals with psychotic disorders. We chose to study individuals with family history of psychosis because they display a hyperdopaminergic phenotype [5], [6] without the confounding treatment of antipsychotic medication, which directly competes with [^11^C]raclopride. Specifically, our hypothesis is that first-degree relatives of individuals with psychotic disorders will show a reduction in adaptive, stimulus-driven dopamine release during fear conditioning. We test if lower dopamine release is associated with measures of subclinical negative or positive symptoms [20]. Finally, in a larger sample that only completed the

Pavlovian fear conditioning fMRI task, we also test whether behavioural ratings of the cues are associated with subclinical negative and positive symptomatology.

## Materials & Methods

### General outline of the study

We employed a visual fear conditioning paradigm during which computer-generated faces were presented to participants [21], [22]. After completing the clinical interview, each participant underwent the fear conditioning task using simultaneous functional magnetic resonance imaging (fMRI) & skin conductance recordings (*N* = 74). A subset of individuals underwent multimodal neuroimaging including a baseline T1-weighted MRI, baseline positron emission tomography (PET) scan and the following day, another T1-weighted MRI, and simultaneous task-based PET, fMRI & skin conductance recordings (*N* = 28). Four runs of fear conditioning were completed in the larger and smaller groups. Baseline and task PET days were counterbalanced and the inter-scan interval was approximately 24 hours for all participants, with the two scans never done on the same day. An explicit recall task was administered no more than 2 hours after acquisition. A second recall task was administered either in-person or virtually approximately 24 hours after acquisition. See ***supplementary methods*** for a full description of the skin conductance recordings, visual fear conditioning paradigm & contingency memory task (**Figure S1**).

We administered [^11^C]raclopride with 2 separate bolus scans is the most established design used to quantify dopamine release in the living human brain [23]. The analysis was restricted to the striatum since [^11^C]raclopride binding is not displaceable elsewhere in the brain [24]. We scanned individuals twice and used a task design that maximized the displacement of the tracer during the task PET session which avoids challenges in modeling bolus + constant infusion [25]. Thus, detection of dopamine release during fear acquisition is maximized with these considerations [26].

### Participants

The study protocol was approved by The Royal Ottawa Healthcare Group Research Ethics Board (REB #: 2019003) and Declaration of Helsinki. All participants gave a written informed consent before entering the study. Control participant status was confirmed using the MINI International Neuropsychiatric Interview for the DSM-5 [27]. Control participants with severe medical illness, head trauma resulting in unconsciousness, current psychiatric illness including substance use disorders (including nicotine), past psychosis, positive urine drug test, and/or contraindications for MRI scanning were excluded from the study. FDRs were recruited in collaboration with psychiatrists at the ROMHC and through print advertisements in the community. FDR status was confirmed using the Family Interview for Genetic Studies - Psychosis Checklist [28] and confirmation of the proband’s diagnosis through electronic medical records when possible. Two FDR participants screened positively for a current mental health disorder (one with social anxiety disorder and the other with generalized anxiety disorder).

Although they screened positively, these two FDRs were included in the study. They were not taking any medication to treat their disorders.

### PET/MR Image acquisition & analysis

PET/MR images were acquired using a 3T mMR Biograph PET-MR system with a 12-channel head coil (Siemens, Erlangen, Germany). The full acquisition parameters and analysis details are described in ***supplementary methods***.

### Statistical analysis

All statistical analyses were conducted in R v4.5.1. Linear mixed effects models and subsequent marginal means were conducted and then extracted using the ‘nlme’ package. Then marginal means were inputted for post-hoc comparisons using the ‘emmeans’ package. Correlations were done using the native R ‘cor’ function. Ordinary least squares (OLS) regressions were done using the native R ‘lm’ function. All post-hoc comparisons were corrected using False Discovery Rate (FDR). All plots were made using seaborn v0.13.2 [29]. All cortical and subcortical visualizations were made using surfplot v0.2.0 [30] and subcortex_visualization v0.1.12 [31].

## Results

### Dopamine is released in the striatum during fear conditioning in humans

Prior to characterizing fear-elicited dopamine release in those with a family history of psychosis, we sought to quantify dopamine release in individuals without a family history.

Sixteen individuals without a family history of psychotic disorders (*N* = 16) completed the Pavlovian fear acquisition task over four runs. Age, sex, injected tracer dose, injected molar activity, motion during the PET scan, US shock intensity, self-reported subclinical symptoms and behavioural fear responses of this control group are presented in **Table 1** and **Table S1**.

**Table 1.** This study includes two groups of participants: a simultaneous PET/fMRI group (*N* = 28) and a second fMRI only group that completed the same task without PET (*N* = 46) that was combined for fMRI analysis (*N* = 74).

| | Simultaneous PET/fMRI ($N = 28$) | | | fMRI ($N = 74$) | | |
| --- | --- | --- | --- | --- | --- | --- |
| | CTR<br>$n = 16$ | FDR<br>$n = 12$ | test | CTR<br>$n = 50$ | FDR<br>$n = 24$ | test |
| **Age**<br>(SD) | 33 (9.8) | 33 (11) | $t_{(26)} = 0.03$,<br>$p = .97$ | 30 (8.5) | 32 (9.0) | $t_{(72)} = -0.71$,<br>$p = .48$ |
| **Sex**<br>(male) | 9/16 (56.3%) | 7/12 (58.3%) | $\chi^2 < 0.01$,<br>$p = 1.0$ | 26/50 (52%) | 12/24 (50%) | $\chi^2 < 0.01$,<br>$p = 1.0$ |
| **US**<br>(SD) [mA] | 2.7 (0.95) | 2.9 (1.04) | $t_{(26)} = 0.48$,<br>$p = .63$ | 2.2 (0.98) | 2.7 (1.0) | $t_{(72)} = 2.0$,<br>$p = .04$ |
| **Baseline dose**<br>(SD) [MBq] | 300 (42) | 310 (19) | $t_{(26)} = 1.04$,<br>$p = .31$ | | | |
| **Task dose**<br>(SD) [MBq] | 280 (31) | 290 (22) | $t_{(26)} = 1.08$,<br>$p = .29$ | | | |
| **Baseline MA**<br>(SD) [mCi/umol] | 1600 (1100) | 1300 (410) | $t_{(26)} = -0.67$,<br>$p = .51$ | | | |
| **Task MA**<br>(SD) [mCi/umol] | 1300 (420) | 1300 (680) | $t_{(26)} = 0.28$,<br>$p = .78$ | | | |
| **Baseline FD**<br>(SD) [mm] | 1.6 (0.64) | 1.5 (0.50) | $t_{(26)} = -0.74$,<br>$p = .46$ | | | |
| **Task FD**<br>(SD) [mm] | 1.6 (0.81) | 1.6 (0.66) | $t_{(26)} < 0.01$,<br>$p = .99$ | | | |

*Note:* Demographics (age and sex), the average of four runs of unconditioned stimulus (US) ankle shock intensity in milliamperes (mA) during the fear conditioning session are reported separately for the simultaneous PET/fMRI and combined fMRI only group. Furthermore, the injected dose of the radioligand in megabecquerels (MBq), the injected molar activity (MA) of the radioligand in millicuries per micromole (mCi/umol) and the average framewise displacement (FD; a metric of motion) in millimetres (mm) during the baseline and fear conditioning sessions, are reported for the combined PET/fMRI group. In the combined PET/MR group ( $N = 28$ , 16 CTRs, 12 FDRs), there were no significant differences between control (CTR) and first-degree relative (FDR) participants in age, sex, US shock intensity, injected baseline and task doses/molar activity, or motion on baseline and task day as indexed by FD. In the larger fMRI-only group ( $N = 74$ , 50 CTRs, 24 FDRs), CTR and FDR participants did not differ significantly in age or sex, but the US shock intensity significantly differed.

Dopamine release is quantified as a percent change in dopamine D_2/3_ receptor binding potential (%ΔD_2_R *BP*_ND_) between the baseline scan and the scan during the Pavlovian fear conditioning paradigm (see *methods* and **Figure S1A**). First, we examine D_2_R *BP*_ND_ in a session by region analysis because striatal-projecting dopamine neurons respond differently to fear versus reward [32]–[35]. We find that the fear conditioning task leads to the release of dopamine in the posterior caudate (*n* = 16, μ = 0.116, *SE* = 0.044, *t*_(75)_ = 2.626, *p* = .011 *p*_FDR_ = .032) and posterior putamen (*n* = 16, μ = 0.112, *SE* = 0.044, *t*_(75)_ = 2.549, *p* = .013 *p*_FDR_ = .032) of the striatum in CTRs (**Figure 1a,b; Table S2**).

**Figure 1.**
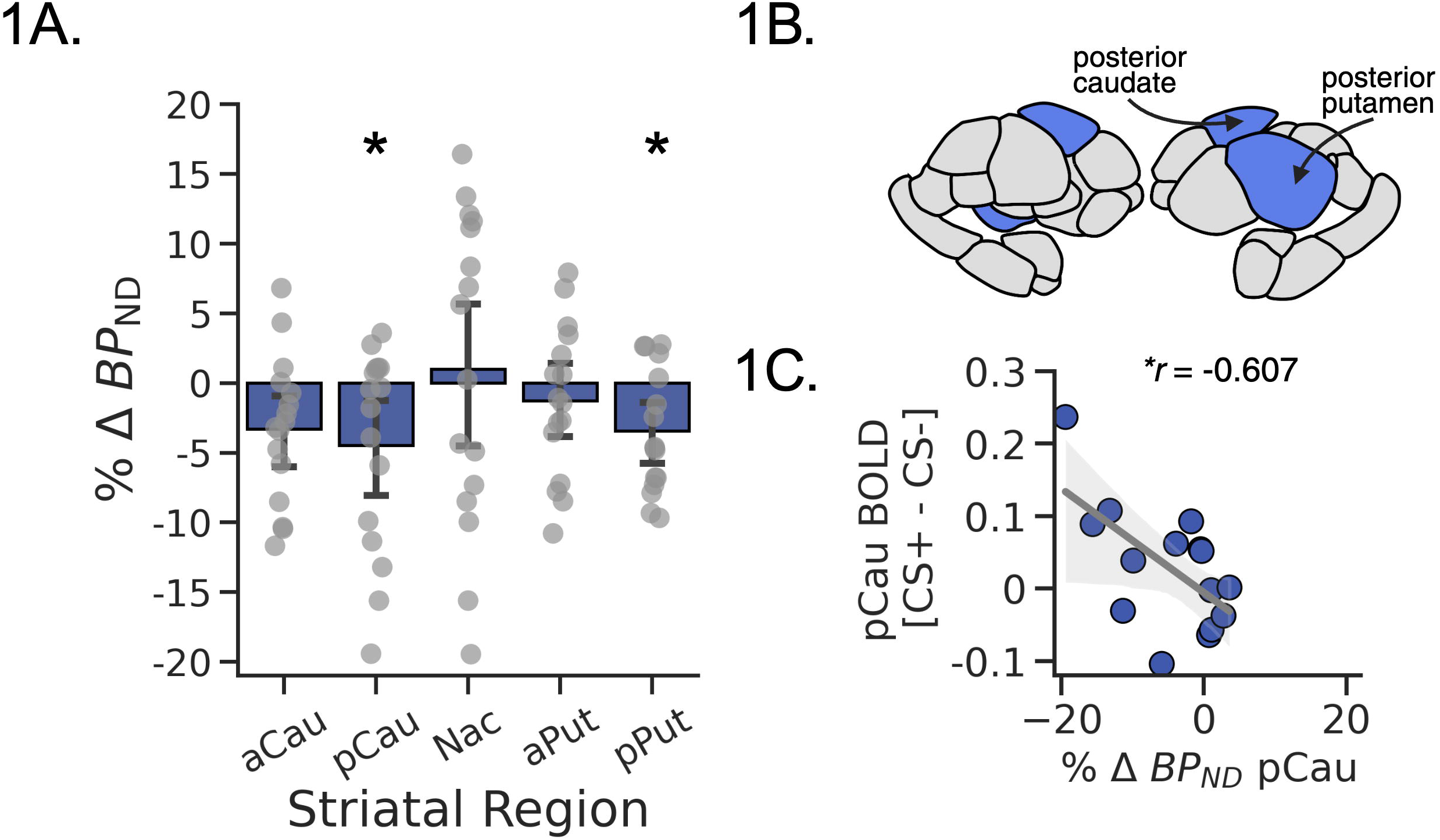
Dopamine is released in the posterior caudate and posterior putamen during fear conditioning in humans. **a.** A linear mixed effects model is used to determine dopamine release. The model included region and session as fixed effects, as well as region and participant as crossed random effects. Post-hoc comparison of regions between sessions reveal D_2_R *BP*_ND_ in the posterior caudate and putamen are significantly decreased during fear conditioning (*n* = 16, μ = 0.112-0.116, *SE* = 0.042, *t*_(75)_ = 2.55-2.62, *p*_FDR_ = .032 [baseline - fear conditioning D_2_R *BP*_ND_]; bar plots with error bars indicate mean %Δ D_2_R*BP*_ND_ ± 95% CI, dot plots represent individual values). **b.** Dopamine is released in the posterior striatal regions (Tian S2 atlas). **c.** Dopamine release in the posterior caudate is associated with greater BOLD signal to the CS+ relative to the CS- (*n* = 16, Pearson’s correlation: *r* = -0.607, *p* = .013, *p*_FDR_ = .025; regression line indicates slope ± 95% CI, scatter plot represents individual values). aCau = anterior caudate; pCau = posterior caudate; Nac = Nucleus accumbens; aPut = anterior putamen, pPut = posterior putamen; %Δ *BP*_ND_ = percent change in binding potential. * = significant *p*-value.

We then examine the link between dopamine release and neural activity. During the fear conditioning two stimuli are presented: the conditioned stimulus (CS+) which is partially reinforced with an unconditioned shock stimulus (US), signalling fear, and a second stimulus that is never reinforced (CS-), signalling safety (see *methods*). We observe an association between %ΔD_2_R *BP*_ND_ and fMRI activation (BOLD response to the contrast of CS+ vs CS-) in the posterior caudate (*n* = 16, *r* = -0.607, *p* = .013, *p*_FDR_ = .025; **Figure 1c**) but not the posterior putamen (*n* = 16, *r* = -0.369, *p* = .159; data not shown). In summary, the largest magnitude of phasic dopamine release occurs in the posterior caudate during fear acquisition and is associated with BOLD activity in the same region.

### Adaptive dopamine release is blunted during fear acquisition in first-degree relatives

To determine if those with an FDR with a psychotic disorder differentially release dopamine during fear acquisition, we compared data from 16 control participants and 12 FDR participants after excluding one FDR for excess motion that compromised the PET kinetic fits (see *methods*) (see **Table 2** for proband information). A linear mixed effects analysis was conducted to examine the interaction between group and striatal region on %ΔD_2_R *BP*_ND_. We find that during fear conditioning, FDR participants release significantly less dopamine in the posterior caudate (*n* = 28, μ = 7.047%, *SE* = 2.762%, *t*_(26)_ = 2.551, *p* = .017, *p*_FDR_ = .034), but not the posterior putamen (*n* = 28, μ = 5.099%, *SE* = 2.762%, *t*_(26)_ = -1.846, *p* = .076) (**Figure 2a,b; Table S3**). To determine if dopamine release is associated with striatal BOLD responses, we also conducted a regression using %ΔD_2_R *BP*_ND_ and group as predictors. We further show that even when controlling for family history of a psychotic disorder, %ΔD_2_R *BP*_ND_ is associated with greater BOLD response to the contrast of CS+ vs CS- (*n* = 28, group: β = -0.067, *SE* = 0.035, *t*_(25)_ = -1.918, *p* = .067; %ΔD_2_R *BP*_ND_: β = -0.005, *SE* = 0.002, *t*_(25)_ = -2.652, *p* = .014) (**Figure 2c**). Importantly, change of participant motion between sessions did not correlate with %ΔD_2_R *BP*_ND_ in the posterior caudate or putamen (N = 28, Pearson’s correlation: all *r* < 0.15, all *p* > 0.44; data not shown). Furthermore, this reduction in dopamine release is not due to between-group differences in US shock intensity, injected dose, baseline D_2_ *BP*_ND_, or motion. Likewise, behavioural learning as indexed by skin conductance recordings during fear acquisition and explicit recall, is not different between FDR and CTR groups of the combined PET/fMRI experiment (**Table 1; Table S1; Table S4**). However, we did find modest differences in fMRI responses in the amygdala and visual association areas in the larger CTR and FDR groups (**Table S8**).

**Figure 2.**
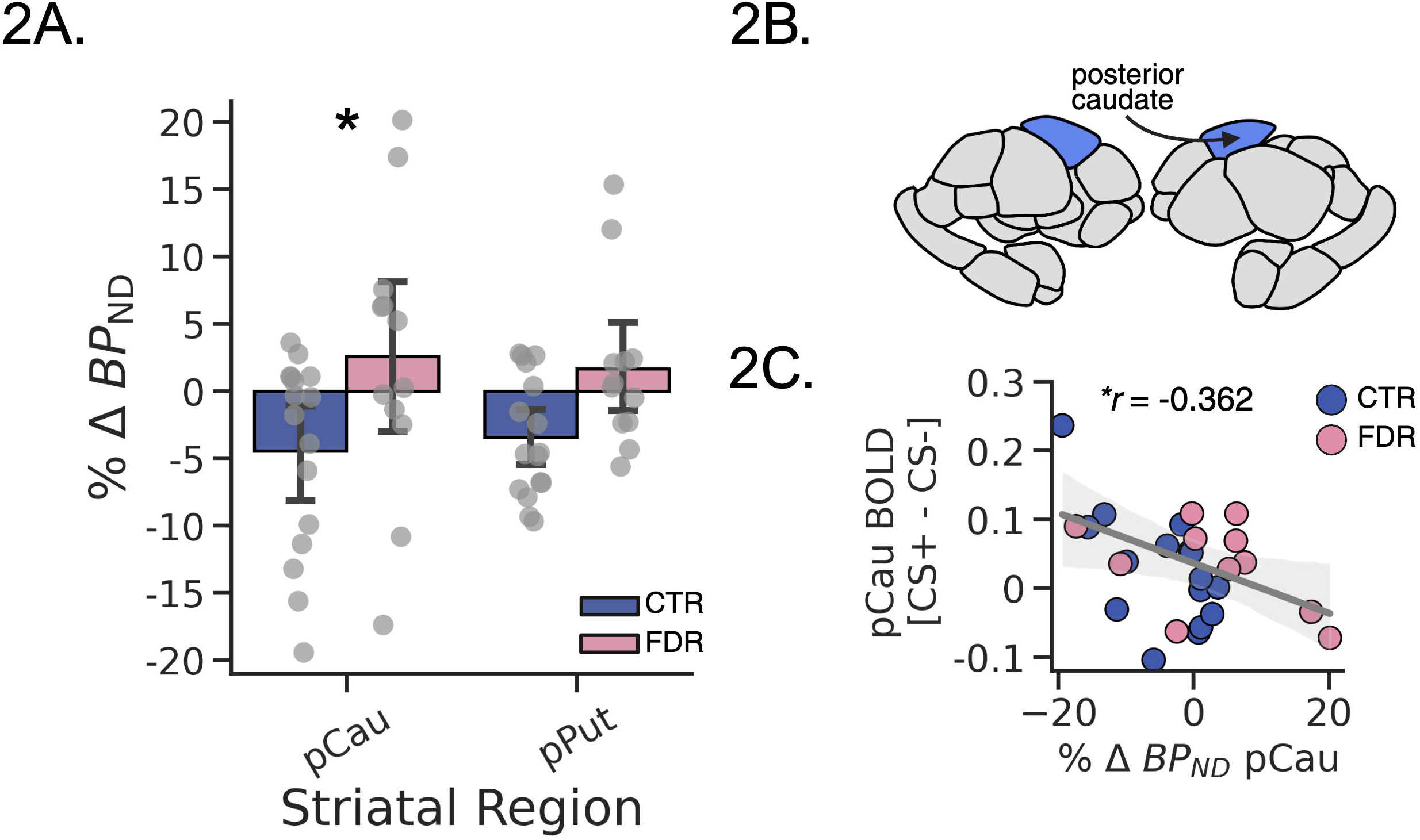
Dopamine release in the posterior caudate is reduced in FDR participants compared to CTRs. **a.** A linear mixed effects model is used to determine differences in dopamine release between FDR and control participants. The model included region and session as fixed effects, and region nested within-participants as a random effect. Post-hoc comparisons indicate that %ΔD_2_R BP_ND_ in the posterior caudate is significantly reduced during fear acquisition (*n* = 28, μ = 7.047%, *SE* = 2.762%, *t*_(26)_ = 2.551, *p* = .017, *p*_FDR_ = .034 [FDR - CTR ΔD_2_R *BP*_ND_]; bar plots with error bars indicate mean %ΔD_2_R *BP*_ND_ ± 95% CI, dot plots represent individual values), **b.** Illustration of the localization of the group difference in the posterior caudate (Tian S2 atlas). **c.** Dopamine release in the posterior caudate is associated with greater BOLD signal to the CS+ relative to the CS- in the whole group when correcting for family history of a psychotic disorder (*n* = 28, regression model, group: β = -0.067, *SE* = 0.035, *t*_(25)_ = -1.918, *p* = .067; %ΔD_2_R *BP*_ND_: β = -0.005, *SE* = 0.002, *t*_(25)_ = -2.652, *p* = .014; regression line indicates slope ± 95% CI, scatter plot represents individual values). pCau = posterior caudate; pPut = posterior putamen; %Δ*BP*_ND_ = percent change in non-displaceable binding potential. * = significant *p*-value.

**Table 2.** Proband relationship status and diagnosis.

|  | **Simultaneous PET/fMRI**<br>**(N = 12)** | **fMRI**<br>**(N = 24)** |
| --- | --- | --- |
| **Relationship** (Sibling/Parent/Both) | 6/6/0 | 10/13/1 |
| **Diagnosis** (SCZ/SA/BP/PNOS) | 7/1/0/3 | 15/5/2/3 |

*Note:* The FDR with both a parent and sibling affected by psychosis contributes 2 individuals to the diagnosis count. SCZ = schizophrenia; SA; schizoaffective disorder; BP; bipolar with psychosis; PNOS; psychosis not otherwise specified.

### Lack of adaptive dopamine release is associated with more paranoid thinking but not with anhedonia

We next examine whether adaptive dopamine release is associated with measures of subclinical positive and negative symptoms, as predicted by the chaotic dopamine hypothesis [19]. Subclinical anhedonia, a negative symptom of schizophrenia, is measured here with the Temporal Experience of Pleasure Scale (TEPS) [36]. We measure subclinical paranoia, a positive symptom, with the Paranoia Checklist [37]. We see that a lack of adaptive dopamine release in the posterior caudate is associated with higher paranoia checklist total scores even when taking family history of psychosis into account (*n* = 28, %ΔD_2_R BP_ND_ β = 0.004, *SE* = 0.002, *t*_(25)_ = 2.074, *p* = .049; group: β = -0.043, *SE* = 0.033, *t*_(25)_ = 1.257, *p* = .22). The association between fear learning and paranoia is driven by higher frequency of self-reported paranoid thoughts, but not conviction or distress subscores (*N* = 28, %ΔD_2_R BP_ND_ β = 0.003, *SE* = 0.001, *t_(25)_* = 2.199, *p* = .037; group: β = -0.008, *SE* = 0.024, *t_(25)_* = -0.334, *p* = .741) (**Figure 3a**) (**Table S9**). In an exploratory analysis, a lack of adaptive dopamine release in the posterior putamen is also associated with a higher paranoia checklist total score (*N* = 28, %ΔD_2_R BP_ND_ β = 0.008, *SE* = 0.003, *t*(25) = 2.806, *p* = .009; group: β = -0.057, *SE* = 0.033, *t*_(25)_ = 1.727, *p* = .097) and a higher frequency score (*N* = 28, regression model: %ΔD2R BP_ND_ β = 0.006, *SE* = 0.002, *t*_(25)_ = 2.866, *p* = .008; group: β = 0.002, *SE* = 0.023, *t*_(25)_ = 0.076, *p* = .940) (**Figure 3b**).

**Figure 3.**
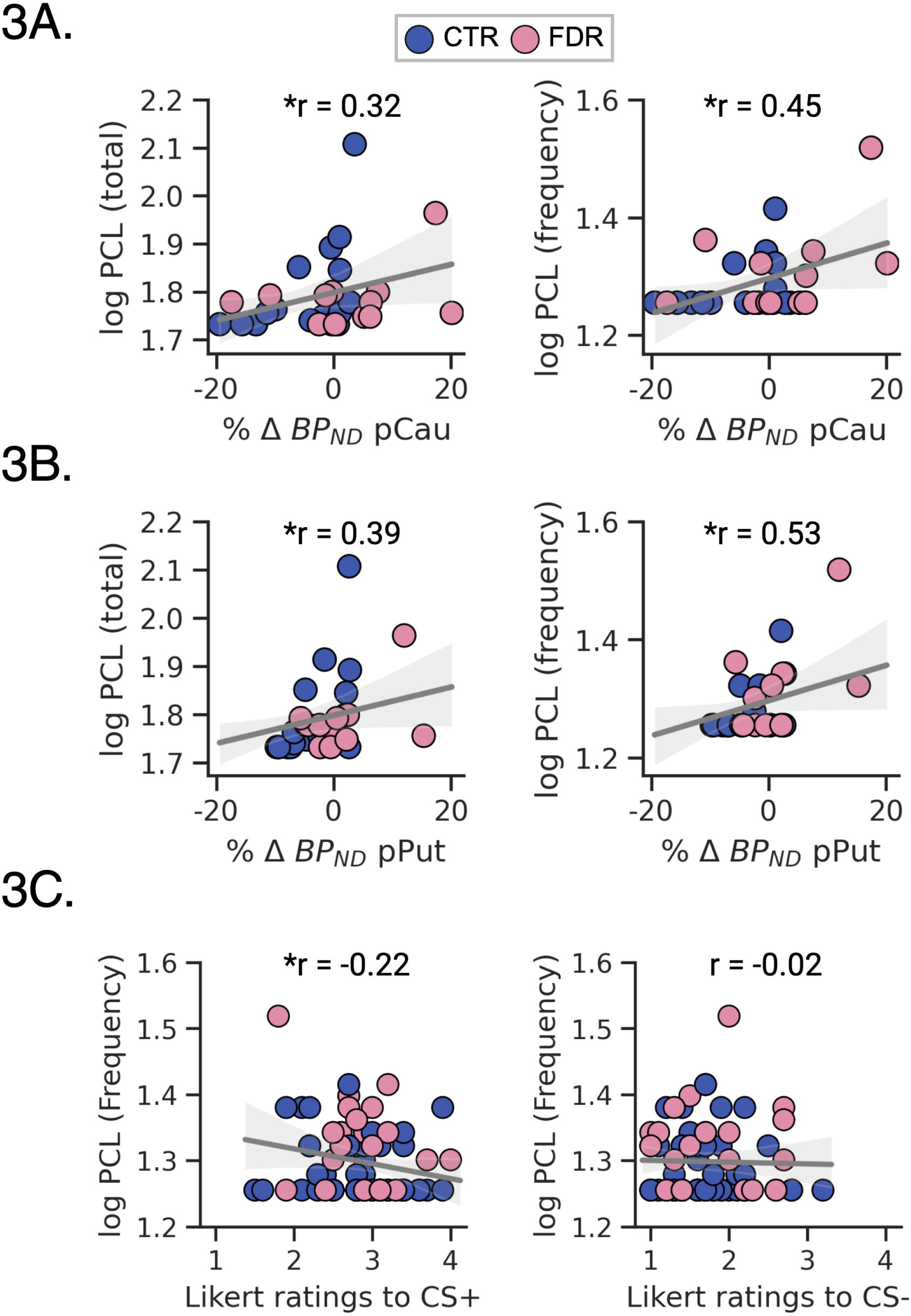
Lack of dopamine release and poor learning about the CS+ predict paranoia checklist frequency scores. **a.** Lack of dopamine release in the posterior caudate is associated with the paranoia checklist total score (*N* = 28, regression model: %ΔD_2_R BP_ND_ β = 0.004, *SE* = 0.002, *t*_(25)_ = 2.074, *p* = 0.049; group: β = -0.043, *SE* = 0.033, *t*(25) = 1.257, *p* = 0.22) and frequency of paranoid thinking (*N* = 28, regression model: %ΔD_2_R BP_ND_ β = 0.003, *SE* = 0.001, *t*_(25)_ = 2.199, *p* = .037; group: β = -0.008, *SE* = 0.024, *t*_(25)_ = -0.334, *p* = .741). **b.** Lack of dopamine release in the posterior putamen is associated with the paranoia checklist total score (*N* = 28, regression model: %ΔD_2_R BP_ND_ β = 0.008, *SE* = 0.003, *t*_(25)_ = 2.806, *p* = .009; group: β = -0.057, *SE* = 0.033, *t*_(25)_ = 1.727, *p* = .097) and frequency of paranoid thinking (*N* = 28, regression model: %ΔD_2_R BP_ND_ β = 0.006, *SE* = 0.002, *t*_(25)_ = 2.866, *p* = .008; group: β = 0.002, *SE* = 0.023, *t*_(25)_ = 0.076, *p* = .940). **c.** Paranoid thinking is predicted by Likert ratings of the CS+ (*n* = 74, regression model for CS+ on day 1: β = -0.024, *SE* = 0.011, *t*_(71)_ = -2.041, *p* = .045) but not Likert ratings of the CS- (*n* = 74, regression model for CS- on day 1: β = -0.003, *SE* = 0.012, *t*_(71)_ = -0.259, *p* = .789). pCAU = posterior caudate; pPUT = posterior putamen; * = significant *p*- value.

This link between fear learning and paranoia is further supported in a larger sample that completed the same task without PET, where lower explicit ratings of the threatening CS+ faces, but not the safe CS- faces, also predicted higher paranoia checklist frequency subscores (*n* = 74, CS+: β = -0.024, *SE* = 0.011, *t*_(71)_ = -2.041, *p* = 0.045; CS-: β = -0.003, *SE* = 0.012, *t*_(71)_ = -0.259, *p* = .789) (**Figure 3c**). In contrast, associations with the TEPS were not supported. That is, we did not find an association between dopamine release in the posterior caudate and self-reported anhedonia, as %ΔD_2_R *BP*_ND_ was unrelated to total, anticipatory, or consummatory subscores on the TEPS (**Table S9**). Altogether, our behavioral results suggest that a lack of adaptive dopamine release during fear learning is associated specifically with the frequency of paranoid thoughts.

### Baseline D_2_Rs but not dopamine release is associated with fear recall

We also examined whether adaptive dopamine release is associated with behavioural indices of fear conditioning. We did not see an association between dopamine release and autonomic responding (skin conductance) or explicit recall (**Table S9**). We also conducted an exploratory analysis between baseline D_2_Rs and these same indices of fear conditioning. Unlike dopamine release, baseline D_2_Rs in the posterior caudate are associated with better recall of the CS+ relative to the CS- (*N* = 28, regression model: D_2_R BP_ND_ β = 0.606, *SE* = 0.218, *t*_(52)_ = 2.777, *p* = 0.008; group: β = 0.083, *SE* = 0.183, *t*_(52)_ = 0.454, *p* = 0.65; day: β = -0.103, *SE* = 0.177, *t*_(52)_ = -0.579, *p* = 0.56). However, baseline D_2_Rs in the posterior caudate are not associated with greater skin conductance towards CS+ relative to CS- (*N* = 28, regression model: D_2_R BP_ND_ β = 0.046, *SE* = 0.071, *t*_(25)_ = 0.648, *p* = 0.523; group: β = 0.067, *SE* = 0.822, *t*_(25)_ = 0.082, *p* = 0.94). In summary, we show that baseline measurements of D_2_Rs in the whole group are associated with better fear discrimination during recall.

## Discussion

Here, we first showed that in humans, fear conditioning releases dopamine in the striatum using [^11^C]raclopride PET. This dopamine release is greatest in the posterior caudate and is proportional to neural activation in the same region measured as BOLD contrast between responses to CS+ and CS- stimuli. Using this paradigm, we then tested the chaotic dopamine hypothesis of schizophrenia. We found that dopamine release is blunted during fear conditioning in people with a family history of a psychotic disorder. The blunted dopamine release was not associated with any differences in fear learning, nor did we find any learning differences between those who had a family history and those who did not. In the whole group, blunted dopamine release was associated with greater paranoia, but not with anhedonia.

Altogether, we provide the first evidence for the impairment in adaptive striatal dopamine release in first-degree relatives of individuals with a psychotic disorder and those who experience subclinical paranoid thoughts.

We report dopamine release in the posterior caudate and putamen during fear acquisition. This finding aligns with rodent studies showing dopamine release in the rodent analogue (striatal tail), a region known to generate a dopamine prediction error that drives fear acquisition, but not recall [34], [35], [38]–[41]. Our finding contrasts with the few available human studies, potentially due to methodological differences like measurement timing (fear acquisition vs. recall), tracer choice, and experimental design [42], [43]. While one study observed indiscriminate striatal dopamine release [43], our findings highlight subregion-specific release within the posterior caudate, supported by correlations between %ΔD_2_R *BP*_ND_ and fMRI activations. This is further supported by a rodent study that reported concurrent dopamine release and striatal neural activity during fear acquisition [41]. In summary, our study is the first to demonstrate dopamine release specifically in the two dorsoposterior striatal areas during fear acquisition in humans.

We find reduced task-based dopamine release in the posterior caudate in a group with a family history of psychotic disorders. There are three previous PET only studies on task-induced dopamine release in psychotic disorders that found a lack of cortical dopamine release during a working memory task and increased nigrostriatal dopamine release during acute nonspecific stress [44]–[46]. However, none of those studies assessed adaptive dopamine release in the striatum. Here, using a simultaneous PET/fMRI design, we present the first *in vivo* evidence of reduced adaptive dopamine release in those with an FDR with a psychotic disorder. We see that the difference in dopamine release between groups is not due to differences in fear learning or engagement on the task, as behavioural indices of learning were not different between those with and without a first degree relative with a psychotic disorder. Given that tonic dopamine release constitutes a small fraction (approximately 0.01%) of the total dopamine released in the synaptic cleft [47], [48], the absence of a detectable change in D_2_ *BP*_ND_ likely indicates a deficiency in adaptive, stimulus-driven dopamine release in response to the CS+, rather than an increase in baseline tonic dopamine release. Furthermore, we found no evidence for baseline D_2_ *BP*_ND_ differences between the two groups. Altogether, these findings support the chaotic dopamine hypothesis by demonstrating a reduction in adaptive dopamine release [19].

On the other hand, we found that a decrease in adaptive dopamine release and poor recall of the CS+ are linked to subclinical, self-reported paranoid thinking, but not to anhedonia. Paranoid thoughts are common in the general population, and paranoia is one of the most heritable symptoms in schizophrenia, often emerging early in the development of psychosis [37], [49]–[52]. Because delusions, like paranoia, are thought to arise from chaotic and out-of-context dopamine release, our finding of reduced adaptive dopamine release correlating with paranoia contrasts with a subcomponent of the chaotic dopamine hypothesis of schizophrenia [19].

Indeed, lack of adaptive dopamine release may lead to the emergence of paranoid delusion, which may help explain why antipsychotics are generally not effective in preventing psychosis in those at clinical high risk [53].

Previous research has shown that tonic increases in dopamine signaling are linked to positive symptoms [54]–[58]. In contrast, reduced adaptive dopamine release, especially towards rewards, is postulated to be related to negative symptoms [19]. A potential pitfall of current dopaminergic theories of schizophrenia (and psychotic disorders) is that they are primarily based on reward learning tasks that typically target the ventral striatum. Yet, it is currently thought that dopaminergic dysregulation in these disorders is more prominent in the dorsal, associative, and sensorimotor striatal areas [11], [59], [60]. Therefore, it is worth noting that the dopamine dysregulation observed in our study is located in the posterior area of the dorsal striatum, an area involved in integrating both sensory and reward prediction errors [61]. Fear conditioning might thus be a valuable paradigm for studying dopaminergic dysregulation in psychotic disorders, potentially contributing to the refinement of existing theories and our understanding of symptom emergence. We propose that inefficiencies in fear learning, possibly mediated by subcortical reductions in phasic dopamine release, might contribute to the development of positive symptoms, like paranoia. One possibility is that a distorted dopamine-reinforced probabilistic map of future risks and rewards might allow less probable explanations to take hold, leading to paranoia [62], [63].

Interestingly, although dopamine release is not associated with behavioural indices of fear learning in the current study, we demonstrate an association with increased baseline D_2_Rs and better fear discrimination at recall, irrespective of FDR status. Indeed, silencing Adora+ (putative D_2_R expressing) tail of striatum neurons increased fear behaviour in the absence of a learned CS [40]. Furthermore, raclopride administration (at pharmacological, not tracer, doses) immediately after fear acquisition reduced fear discrimination at recall [64]. Together, these preclinical studies suggest D_2_Rs in the posterior striatum are essential for computing the appropriate behavioural response towards a CS. Our finding further strengthens this view by demonstrating an association between D_2_R availability in the living human brain and fear discrimination recall, both a few hours after fear acquisition and one day later.

This study is not without limitations. First, despite including 56 [^11^C]raclorpide PET/fMRI scans, our sample size remains modest. Associations between brain activations and behaviour are notoriously difficult to detect and this might explain why we did not detect a link between dopamine release and other fear-related measures [65]. Second, we studied first-degree relatives of individuals with schizophrenia and psychotic disorders and not those individuals themselves. While evidence suggests that the dopamine system is similarly affected in first-degree relatives, future studies should still attempt to test the chaotic dopamine hypothesis in unmedicated individuals with psychotic disorders.

In summary, we show dopamine release in the tail of the striatum of humans during discriminative fear acquisition, and a reduction of this release in the first-degree relatives of those with psychotic disorders. This deficient release is not explained by group differences in fear learning; instead, dopamine release is correlated inversely with self-reported paranoia but not anhedonia. Poor performance on the explicit recall task, reflected by lower Likert ratings towards the shock-paired stimulus (CS+), also correlates inversely with self-reported paranoid thinking. We believe these findings support the hypothesis that lack of stimulus-driven dopamine release, here observed subcortically, may be a component of schizophrenia and psychotic disorders. In conclusion, fear conditioning emerges as a valuable paradigm for investigating dopamine dysregulation in psychosis, offering avenues for refining existing theories and providing a deeper understanding of how psychotic symptoms arise.

## Supporting information

Full Supplement

## Data Availability Statement

Anonymized data is available upon request. Code for the project including aggregate data, analysis and figures are available at https://github.com/rami-hamati/avlraclo.

## Funding

This research was supported by the Emerging Research Innovators in Mental Health Award (The Royal) and a NARSAD grant (The Brain & Behavior Research Foundation).

## Acknowledgements

We would like to acknowledge the participants of the study, especially the first-degree relatives of patients with psychotic illness. Also, thanks to Drs. Pierre Blier, Jeanne Talbot, Georg Northoff and Timothy Lau for providing medical supervision during the positron emission tomography sessions. Also, thanks to the Brain Imaging Centre at The Royal for facilitating this study. Finally, we would like to thank the anonymous donors who provided funding for the Emerging Researchers in Mental Health Award, and ultimately, this study. An oral presentation and conference proceeding related to this data was previously published in *Biological Psychiatry:* https://doi.org/10.1016/j.biopsych.2025.02.236.

## Author Contributions

**RH:** Investigation, Data Curation, Writing - Original Draft, Writing - Review & Editing, Formal Analysis, Visualization. **CS:** Investigation, Data Curation; Writing - Review & Editing. **BC:** Investigation, Data Curation. **HB:** Investigation, Writing - Review & Editing. **KD:** Writing - Review & Editing. **DJH:** Writing - Review & Editing. **CMC:** Methodology, Supervision, Writing - Review & Editing, Funding acquisition. **LT:** Conceptualization, Methodology, Software, Supervision, Writing - Original Draft, Writing - Review & Editing, Funding acquisition.

## Competing Interests

**RH** previously owned shares in Karuna Therapeutics Inc. and received two knowledge dissemination grants from Otsuka Canada Pharmaceutical Inc. to organize local trainee conferences. **LT** received one knowledge dissemination grant from Otsuka Canada Pharmaceutical Inc. & AbbVie Inc. to organize a pan-Canadian schizophrenia conference. The remaining authors have nothing to disclose.

## Notes

https://github.com/rami-hamati/avlraclo

