## Supplementary material for "Blunted fear-evoked striatal dopamine release in individuals with a family history of psychosis": Full Supplement

### Supplementary Methods

#### *Visual fear conditioning paradigm*

The current paradigm consisted of a fear acquisition session, followed by two fear recall sessions [1]. The acquisition phase consisted of four runs (1932 s total; 483 s/ea) during which the CS+ and CS– stimuli were presented 20 and 14 times each, respectively, (34 total presentations; 4 s/ea) in pseudorandom order with a jittered inter-trial interval (range = 6–15 seconds). The CS+ was followed by the unconditioned stimulus (US; an electric shock to the ankle) in 30% of the trials (6/20 presentations), whereas the CS– was never followed by a shock. Thus, participants observed an equal number of nonreinforced CS+ and CS- stimuli in each run. Participants were told that each face stimulus may or may not be followed by an electrical shock, and were asked to recall which faces were followed by a shock after the experiment. The US (0.5 seconds in duration) was generated and delivered using a Coulbourne electric stimulator (Coulbourne Instruments, Allentown, PA) via a cable attached to the left ankle, 0.1 s prior to removal of the CS+. About 30 minutes prior to the scan, the intensity of the US was set by each participant at a level that was "highly annoying but not painful". Afterwards, participants were asked to recall the faces associated with the shock (CS+) approximately 2 and 24 hours after the fear conditioning session.

#### *Skin conductance recordings (SCR)*

SCR were acquired during the fear conditioning task using two MRI-compatible electrodes placed on the participant's left palm. SCR were measured using a BIOPAC MP160 acquisition system (BIOPAC Systems Inc, Goleta, CA) equipped with an EDA100C MRI amplifier. The SCR were collected at a gain of 5 µS/V, using a 1 Hz low pass filter during acquisition, then digitised at 200 Hz. The data were further filtered with a 3 Hz finite impulse response low pass filter, using the AcqKnowledge software package (BIOPAC Systems Inc, Goleta, CA). Stimulus presentations of CS+US, CS+ and CS- were time-stamped onto the SCR recordings using the LabJack u3-LV (LabJack Corporation, Lakewood, CO) digital-to-analogue converter.

#### *Explicit recall task examining contingency memory*

At the end of the fear conditioning task, participants were transferred into a separate procedure room where they completed an explicit recall task immediately after the experiment, no more than two hours after the last fear conditioning run. Participants were shown the four CS+, four CS-, and two novel faces and were asked to rate faces two times per face, in random order (a total of 20 ratings). For each face, participants were asked the following question: "How likely was this face paired with a shock?". Participants rated the faces using a 4-point Likert scale; 1 = "very unlikely", 2 = "somewhat unlikely", 3 = "somewhat likely", 4 = "very likely". Participants repeated the same explicit recall task approximately 24 hours after fear acquisition.

#### *MR Image acquisition*

Images were acquired using a 3T mMR Biograph PET-MR system with a 12-channel head coil (Siemens, Erlangen, Germany). T1-weighted images were acquired using an multiecho MPRAGE sequence [2]; spatial resolution = 1 mm isotropic; matrix = 256 x 256, number of slices = 192, repetition time (TR) = 2500 ms, echo time (TE) = 1.69, 3.55, 5.41, and 7.27 ms, flip angle = 7 degrees. T2*-weighted (functional MRI) images were acquired during the fear conditioning task; spatial resolution = [3 x 3 x 3.21 mm], matrix = 64 x 64, number of slices = 210, TR = 2300 ms, TE = 28 ms, flip angle = 80 degrees.

#### *MR preprocessing & analysis*

Results included in this manuscript come from preprocessing performed using fMRIPrep 24.0.0 ([3], [4]; RRID:SCR_016216), which is based on Nipype 1.8.6 ([5]; RRID:SCR_002502). To note, sessions one and two were analysed separately, and the boilerplate was modified to include the correct number of scans. All parameters are exactly the same between sessions. Fully preprocessed BOLD images were post-processed using Nilearn 0.11.1 ([6]; RRID:SCR_001362) in python 3.11.9.

#### *Anatomical data preprocessing*

A total of two T1-weighted (T1w) (one T1w ses-001; one T1w ses-002) images are found within the input BIDS dataset. The T1w image is corrected for intensity non-uniformity (INU) with N4BiasFieldCorrection [7], distributed with ANTs 2.5.1 ([8], RRID:SCR_004757), and used as T1w-reference throughout the workflow. The T1w-reference is then skull-stripped with a Nipype implementation of the antsBrainExtraction.sh workflow (from ANTs), using OASIS30ANTs as target template. Brain tissue segmentation of cerebrospinal fluid (CSF), white-matter (WM) and gray-matter (GM) is performed on the brain-extracted T1w using fast (FSL (version unknown), [9]; RRID:SCR_002823). Brain surfaces are reconstructed using recon-all (FreeSurfer 7.3.2, [10]; RRID:SCR_001847), and the brain mask previously estimated is refined with a custom variation of the method to reconcile ANTs-derived and FreeSurfer-derived segmentations of the cortical gray-matter of Mindboggle ([11]; RRID:SCR_002438). Volume-based spatial normalization to two standard spaces (MNI152NLin6Asym, MNI152NLin2009cAsym) is performed through nonlinear registration with antsRegistration (ANTs 2.5.1), using brain-extracted versions of both T1w reference and the T1w template. The following templates are were selected for spatial normalization and accessed with TemplateFlow (24.2.0, [12]): FSL's MNI ICBM 152 non-linear 6th Generation Asymmetric Average Brain Stereotaxic Registration Model [[13], RRID:SCR_002823; TemplateFlow ID: MNI152NLin6Asym], ICBM 152 Nonlinear Asymmetrical template version 2009c [[14], RRID:SCR_008796; TemplateFlow ID: MNI152NLin2009cAsym].

#### *Functional data preprocessing*

For each of the eight BOLD runs (three resting-state fMRI ses-001; four task-based fMRI ses-002; one resting state fMRI ses-002) found per participant (across all tasks and sessions), the following preprocessing is performed. First, a reference volume is generated, using a custom methodology of fMRIPrep, for use in head motion correction. Head-motion parameters with respect to the BOLD reference (transformation matrices, and six corresponding rotation and translation parameters) are estimated before any spatiotemporal filtering using mcflirt (FSL, [15]). The BOLD reference is then co-registered to the T1w reference using bbregister (FreeSurfer) which implements boundary-based registration [16]. Co-registration is configured with six degrees of freedom. Several confounding time-series are calculated based on the preprocessed BOLD: framewise displacement (FD), DVARS and three region-wise global signals. FD is computed using two formulations following Power (absolute sum of relative motions [17]) and Jenkinson (relative root mean square displacement between affines [15]). FD and DVARS are calculated for each functional run, both using their implementations in Nipype (following the definitions by [18]. The three global signals are extracted within the CSF, the WM, and the whole-brain masks. Additionally, a set of physiological regressors are extracted to allow for component-based noise correction (CompCor, [19]). Principal components are estimated after high-pass filtering the preprocessed BOLD time-series (using a discrete cosine filter with 128s cut-off) for the two CompCor variants: temporal (tCompCor) and anatomical (aCompCor). tCompCor components are then calculated from the top 2% variable voxels within the brain mask. For aCompCor, three probabilistic masks (CSF, WM and combined CSF+WM) are generated in anatomical space. The implementation differs from that of Behzadi et al. in that instead of eroding the masks by 2 pixels on BOLD space, a mask of pixels that likely contain a volume fraction of GM is subtracted from the aCompCor masks. This mask is obtained by dilating a GM mask extracted from the FreeSurfer's aseg segmentation, and it ensures components are not extracted from voxels containing a minimal fraction of GM. Finally, these masks are resampled into BOLD space and binarized by thresholding at 0.99 (as in the original implementation). Components are also calculated separately within the WM and CSF masks. For each CompCor decomposition, the k components with the largest singular values are retained, such that the retained components' time series are sufficient to explain 50% of variance across the nuisance mask (CSF, WM, combined, or temporal). The remaining components are dropped from consideration. The head-motion estimates calculated in the correction step are also placed within the corresponding confounds file. The confounding time series derived from head motion estimates and global signals are expanded with the inclusion of temporal derivatives and quadratic terms for each [20]. Frames that exceeded a threshold of 0.5 mm FD or 1.5 standardized DVARS are annotated as motion outliers. Additional nuisance time series are calculated by means of principal components analysis of the signal found within a thin band (crown) of voxels around the edge of the brain [21]. The BOLD time-series are resampled onto the following surfaces (FreeSurfer reconstruction nomenclature): fsaverage5. All resamplings can be performed with a single interpolation step by composing all the pertinent transformations (i.e. head-motion transform matrices, susceptibility distortion correction when available, and co-registrations to anatomical and output spaces). Gridded (volumetric) resamplings are performed using nitransforms, configured with cubic B-spline interpolation. Non-gridded (surface) resamplings are performed using mri_vol2surf (FreeSurfer).

Many internal operations of fMRIPrep use Nilearn 0.10.4 ([6]; RRID:SCR_001362), mostly within the functional processing workflow. For more details of the pipeline, see the section corresponding to workflows in fMRIPrep's documentation.

#### *Task-based fMRI analysis*

Fully preprocessed BOLD images were post-processed using Nilearn 0.11.1 ([6]; RRID:SCR_001362) in python 3.11.9. Briefly, a first-level model was constructed with CS+, CS-, and CS+ shock events and a 'simple' confound strategy to fit a glover haemodynamic response function [22]. Afterwards, effect size contrast maps were computed per run, per participant: fear contrast (CS+ versus CS-). A total of four contrast images were computed, then averaged together to generate a single contrast map per participant. Then, the Glasser [23] and Tian atlases [24] were used to extract contrast beta values per parcel (196 parcels; 180 cortical and 16 subcortical parcels bilaterally).

#### [11C]raclopride PET acquisition

PET images were acquired once during baseline and once during the fear conditioning task; spatial resolution = [4.17 x 4.17 x 2.03 mm]; target injected dose = 300 MBq of [$^{11}$C]raclopride (bolus), number of sinograms = 2540, scan time = 60 minutes. All 58 [$^{11}$C]raclopride injections ($N$ = 29) were administered between 11:00 and 15:00 (one scan per participant per day); 23 of 29 completed experiments are no more than one day apart, four experiments were completed two days apart and two experiments were completed eight to nine days apart.

##### *PET preprocessing*

For each of two 60-minute dynamic PET runs found per participant (across scanning sessions), the following preprocessing was performed. Attenuation correction was calculated using the pseudoCT method, which is an MR-based (T1w) technique to generate μ-maps [25]. Subsequently, PET images were reconstructed with the μ-map applying 3 mm Gaussian smoothing, grouped into 1-minute bins (60 frames per image) to allow for minute-by-minute motion correction. For the remainder of the PET preprocessing analysis, the PETsurfer pipeline was used [26], [27]. First, head motion correction [FreeSurfer mc-afni2] was applied to the reconstructed PET image, and the resulting motion-corrected image was used to generate a summed template volume for co-registration to the T1w [FreeSurfer mri_concat; [28]]. The PET template image was then co-registered to the T1w reference [FreeSurfer mri_coreg] and the resulting transformation matrix was applied to the PET image [FreeSurfer mri_vol2vol]. Finally, the T1w reference was co-registered to MNI152nlin6asym space [FreeSurfer mni152reg] and the transformation matrix was applied to the PET image in T1w space [FreeSurfer mri_vol2vol]. The resulting image is a PET scan in MNI152nlin6asym space with 60 motion-corrected frames.

##### *PET analysis*

A total of 58 [$^{11}$C]raclopride scans were collected ($N$ = 29). There was no difference in the injected dose (paired-samples t-test: $t_{(28)}$ = 2.034, $p$ = .0515) or injected molar activity (paired-samples t-test: $t_{(28)}$ = 0.8575, $p$ = .3985) between the baseline session (dose: $\mu$ = 302.76 MBq, $SD$ = 33.63; molar activity: $\mu$ = 1447.5 mCi/umol, $SD$ = 839.64) and task session (dose: $\mu$ = 286.99 MBq, SD = 27.31; molar activity: $\mu$ = 1299.9 mCi/umol, $SD$ = 526.89). $D_2R$ $BP_{ND}$ was quantified with multi-linear reference tissue model 2 (MRTM2) [29] using a custom FreeSurfer/kinfitr pipeline [ROI-wise] [26], [27], [30]. $D_2R$ $BP_{ND}$ is defined as the concentration of target protein ($D_{2/3}$ receptors) in the specifically bound compartment (striatum), relative to the non-displaceable compartment (cerebellum). Striatal ROIs were delineated using an unbiased striatal atlas [24] using the S2 level of granularity (each ROI >298 voxels) and the cerebellar ROI was delineated using the left and right cerebellar cortex ROIs from the FreeSurfer atlas in MNI152 space, and then eroded (fslmaths ero). Then, average time-activity curves were generated for each striatal ROI (FreeSurfer mri_binarize & mri_segstats). The time-activity curves were inputted to the software package kinfitr for the kinetic modeling of $D_2R$ $BP_{ND}$ [30]. Because we do a minute-by-minute motion correction, we only excluded participants that produced less than excellent goodness-of-fit in the kinetic modelling (less than $R^2 < 0.80$). This resulted in one exclusion, and a total of 28 individuals for the final analysis (16 CTR/12 FDR).

Importantly, participant motion did not differ between baseline and fear conditioning sessions (*N* = 28, paired-samples t-test: $t_{(27)} = -0.691$, p = 0.495). MRTM2 goodness-of-fits using the Tian S2 atlas remained above 0.8 (range of $R^2 = 0.84 – 0.99$) only when the nucleus accumbens shell and core were combined into a single ROI; otherwise, $R^2$ values failed to reach this standard. Thus, to balance accurate $BP_{ND}$ estimates and retain granularity, all of our primary analyses use a total of five striatal ROIs: nucleus accumbens, anterior caudate, anterior putamen, posterior caudate and posterior putamen. Finally, $\%\Delta D_2R$ $BP_{ND,}$ our index of dopamine release, was calculated using the following equation:

$$\%\Delta BP_{ND} = [\frac{(BP_{ND}task - BP_{ND}baseline)}{BP_{ND}baseline}] \times 100$$

### Supplementary Results

#### The visual fear discrimination paradigm elicits fear responses but these do not differ between groups

Before examining differences in fear learning between controls and FDRs, the expected learning in the combined PET/fMRI group (*N* = 28) and the larger fMRI group (*N* = 74) are first confirmed. Here, we examine if the fear conditioning task elicited greater skin conductance responses to the CS+, correct recall of the CS+ and typical fMRI responses during fear conditioning across all four runs of fear acquisition using linear mixed effect models. Indeed, expected skin conductance responses, Likert ratings during explicit recall and fMRI responses towards the CS+ are confirmed (**Figure S2; Tables S5, S6**). Reaction times during explicit ratings did not differ between CS+ and CS- (**Figure S2; Table S7**). Next, we find no between-group differences in skin conductance recordings and the explicit recall task (**Tables S5, S6, S7**). Thus, differences in behavioural indices of fear expression between groups do not explain differences in dopamine release between the groups. Finally, we examine between-group differences in the CS+ versus CS- fMRI BOLD responses using the Glasser atlas for the extraction of cortical ROIs and the Tian S2 atlas for the extraction of subcortical ROIs. An exploratory linear mixed effects model reveals several higher order visual and associative areas are differentially activated in controls versus FDRs, as well as the medial amygdala (**Table S8**). Activity in significantly different regions between-groups are not predicted by dopamine release in the posterior caudate (**Table S8**). Thus, BOLD differences between groups do not explain the difference in dopamine release between groups.

### Supplementary Figures

#### Visual depiction of the experimental design

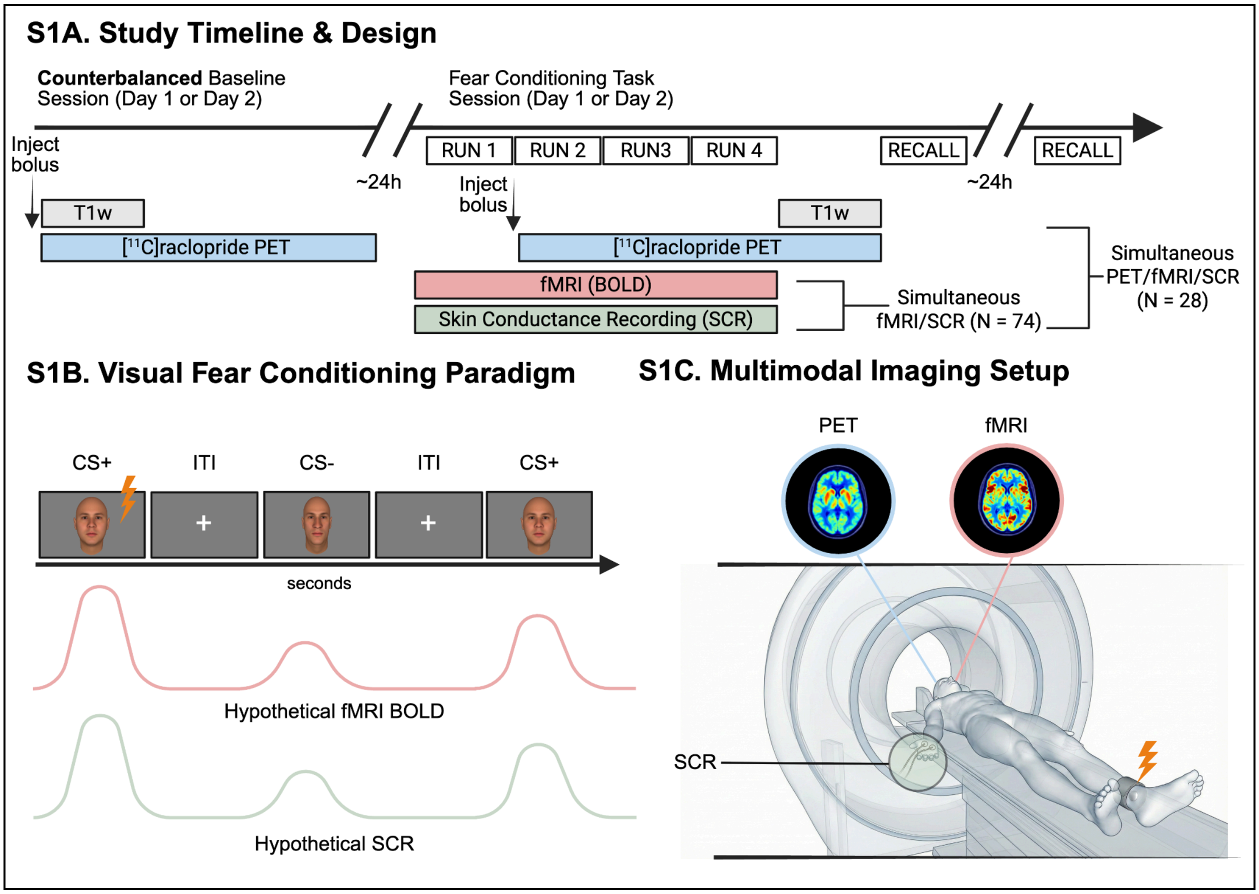


**Figure S1.** Visual depiction of the experimental design and fear conditioning task. **A.** A subset of 28 individuals completed the combined PET/fMRI arm. Here, participants completed one 60-minute [$^{11}$C]raclopride PET scan during baseline. Approximately one day later, participants are brought back for a fear acquisition scan. Participants complete four runs of a visual fear conditioning paradigm in the MRI scanner, with simultaneous acquisitions of fMRI and SCR. [$^{11}$C]raclopride is injected immediately after termination of the first fear conditioning run, and then participants complete three more runs. Afterwards, participants complete an explicit recall of the visual pairings no more than two hours after the last fear conditioning run. Finally, participants complete a second explicit recall approximately one day after the last fear conditioning run. These 28 individuals that completed the combined PET/fMRI arm are part of a larger group (*N* = 74) that completed simultaneous fMRI and SCR, then recall. **B.** The top panel depicts an example of a single fear conditioning run. Participants are presented with a new pair of pseudorandomized faces at each run to ensure continuous learning throughout the scan. One run of fear acquisition consisted of 20 CS+ presentations (6 paired with a shock, ~33% reinforcement rate, 4 seconds each), 14 CS- presentations (4 seconds each) and a jittered inter-trial interval (6-15 seconds each) for a total of 483 seconds (see *methods* for more details).

The bottom panel represents theoretical blood-oxygen-dependent-level (BOLD) and SCR signals. **C.** An example illustration of an individual undergoing simultaneous PET/fMRI/SCR.

#### The visual fear discrimination paradigm elicits fear responses but these do not differ between groups

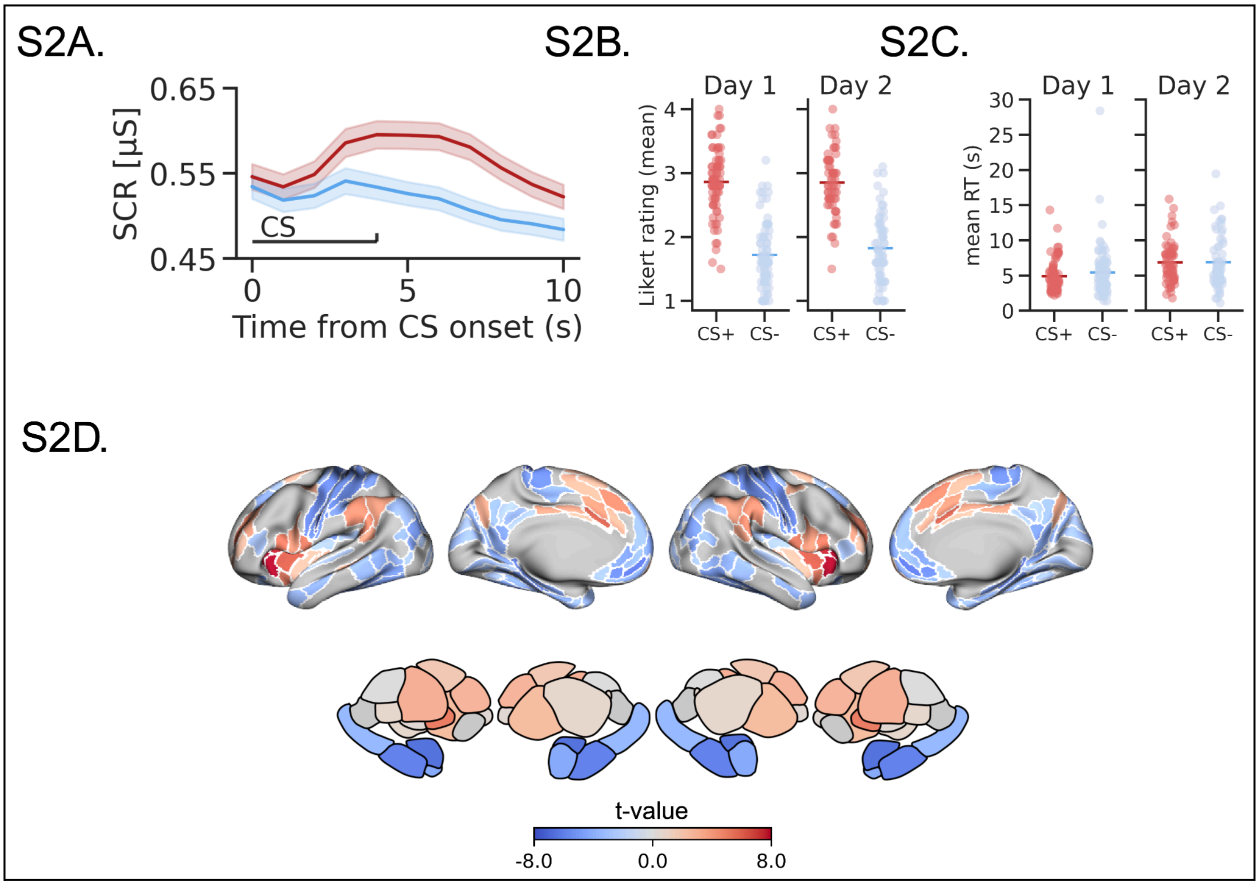


**Figure S2.** Area Under the Curve (AUC) of the deconvolved phasic skin conductance response from 0-10 s (measured in microsiemens [µS]), explicit recall scores representing contingency memory and BOLD responses confirm learning during the visual fear discrimination paradigm. **a.** A linear mixed effects model reveals the CS+ (red) elevated skin conductance responses relative to the CS- (blue). Examination of the main effects of the model using ANOVA reveals a significant effect of CS type, but not group, in the combined PET/fMRI (*N* = 28, $F_{(1,111)} = 6.404$, $p = .0128$) and the larger fMRI group (*N* = 69, $F_{(1,275)} = 12.02$, $p = .0006$). **b.** A linear mixed effects model reveals that participants rated the CS+ (red) as more likely to be paired with the shock relative to the CS- (blue), 2 hours after the last fear conditioning run and the next day. Examination of the main effects of the model using ANOVA reveals a significant effect of CS type, but not group or day, in the combined PET/fMRI (*N* = 28, $F_{(1,334)} = 246.1$, $p < .0001$) and the larger fMRI group (*N* = 74, $F_{(1,782)} = 578.4$, $p < .0001$) **c.** However, a linear mixed effects model did not reveal differences in reaction time (RT) when rating CS+ and CS- stimuli when examining main effects in the combined PET/fMRI (*N* = 28, $F_{(1,334)} = 0.207$, $p = .6491$) and the larger fMRI group (*N* = 74, $F_{(1,782)} = 1.452$, $p = .2285$). **d.** A one-sample *t*-test of BOLD activity during the fear conditioning task indicates that 38 parcels exhibit BOLD activity favoring the CS+ relative to CS- (red), while 75 parcels show BOLD activity favoring the CS- relative to CS+

(blue) ($N$ = 74, all $t_{(73)} > |2.2|$, all $p < 0.05$; fdr-adjusted for 196 parcels). Importantly, we find similar changes in BOLD activity in these regions as previous meta-analyses of fear conditioning [31], [32].

### Supplementary Tables

**Table S1.** Temporal Experience of Pleasure Scale (TEPS), State Paranoia Checklist (paranoia checklist), explicit recall, and skin conductance from the combined PET/fMRI group ($N$ = 28).

| | Controls (N = 16) | FDRs (N = 12) | Test | Statistic |
|---|---|---|---|---|
| **TEPS** | | | | |
| Anticipatory Subscore \| Mean (SD) | 42 (8.0) | 43 (5.5) | t-test | t(26) = -0.22, p = 0.83 |
| Consummatory Subscore \| Mean (SD) | 39 (6.4) | 38 (5.4) | t-test | t(26) = 0.42, p = 0.68 |
| Total Score \| Mean (SD) | 81 (13) | 81 (9.4) | t-test | t(26) = 0.08, p = 0.94 |
| **paranoia checklist** | | | | |
| Frequency Subscore \| Median [Min, Max] | 18 [18,26] | 19 [18,33] | wilcoxon | W = 75.5, p = 0.29 |
| Conviction Subscore \| Median [Min, Max] | 20 [18,73] | 21 [18, 34] | wilcoxon | W = 89, p = 0.76 |
| Distress Subscore \| Median [Min, Max] | 19 [18, 37] | 18 [18, 25] | wilcoxon | W = 112.5, p = 0.42 |
| Total Score \| Median [Min, Max] | 57 [54, 130] | 60 [54,92] | wilcoxon | W = 86, p = 0.66 |
| **Explicit Recall** | | | | |
| *Day 1* | | | | |
| CS+ \| Mean (SD) | 3.22 (0.96) | 3.03 (0.98) | Linear mixed effect model | F(1,111) = 122, p < 0.0001 **[CS- vs. CS+]** F(1,26) = 0.03, p = 0.86 **[group]** |
| CS- \| Mean (SD) | 1.73 (1.00) | 1.86 (1.02) | | |
| *Day 2* | | | | |
| CS+ \| Mean (SD) | 3.17 (0.94) | 3.07 (1.07) | Linear mixed effect model | F(1,111) = 91.1, p < 0.0001 **[CS- vs. CS+]** F(1,25) = 0.15, p = 0.69 **[group]** |
| CS- \| Mean (SD) | 1.90 (1.00) | 1.86 (1.07) | | |
| **Skin conductance** | | | | |
| CS+ \| Mean (SD) | 6.81 (7.49) | 9.74 (12.3) | Linear mixed effect model | F(1,111) = 6.40, p = 0.013 **[CS- vs. CS+]** F(1,26) = 0.75, |
| CS- \| Mean (SD) | 6.21 (6.77) | 8.74 (10.1) | | |

| | | | | |
|---|---|---|---|---|
| | | | | p = 0.39<br>**[group]** |

**Table S2.** %$\Delta D_2R$ $BP_{ND}$ data are fit using a linear fixed effects model in 16 control participants x 2 timepoints x 5 regions = 150 total observations. As per the equation below [lme(bpnd ~ session*region, random = list(*participant_*id = ~ 1, region = ~ 1), data=filtered_data, method="REML")], we specify fixed effects of *region* and *session*. We also specify two random effects of *participant_id* and *region.* This syntax specifies *1|participant_id* to model both overall individual differences as well as *1|region in participant_id* to model region-specific individual differences. Then, the *emmeans* package in R is used to extract marginal means from the model. The results of the pairwise comparisons [BL - FC] are presented below. SE = standard error; df = degrees of freedom.

| region | estimate | SE | df | t.ratio | p.value | fdr.adj.p |
|---|---|---|---|---|---|---|
| Nucleus accumbens | -0.017 | 0.044 | 75 | -0.378 | 0.7064 | 0.706 |
| Anterior caudate | 0.084 | 0.044 | 75 | 1.916 | 0.0592 | 0.099 |
| Anterior putamen | 0.038 | 0.044 | 75 | 0.871 | 0.3866 | 0.483 |
| Posterior caudate | 0.116 | 0.044 | 75 | 2.626 | 0.0105 | 0.032 |
| Posterior putamen | 0.115 | 0.044 | 75 | 2.549 | 0.0129 | 0.032 |

**Table S3.** %$\Delta D_2R$ $BP_{ND}$ data are fit using a linear fixed effects model in 16 control participants + 12 FDR participants. One difference value was obtained from each participant x 2 regions = 54 total observations. As per the equation [*lme(bpnd ~ region*group, random = list(participant_id = ~ 1, region = ~ 1), data=filtered_data, method="REML")*], we specify fixed effects of *region* and *session*. We also specify two random effects of *participant_id* and *region.* This syntax specifies *1|participant_id* to model both overall individual differences as well as *1|region in participant_id* to model region-specific individual differences. Then, the *emmeans* package in R is used to extract marginal means from the model. The results of the pairwise comparisons [FDR - CTR] are presented below. SE = standard error; df = degrees of freedom.

| region | estimate | SE | df | t.ratio | p.value | fdr.adj.p |
|---|---|---|---|---|---|---|
| Posterior caudate | 7.047 | 2.762 | 26 | 2.551 | 0.017 | 0.034 |
| Posterior putamen | 5.100 | 2.762 | 26 | 1.846 | 0.076 | 0.076 |

**Table S4.** Baseline $D_2R$ $BP_{ND}$ data are fit using a linear fixed effects model in 16 control participants + 12 FDR participants. One $BP_{ND}$ value is obtained from each participant x 2 regions = 54 total observations. As per the equation *[lme(bpnd ~ group*region, random = list(participant_id = ~ 1, region = ~ 1), data=filtered_data, method="REML")*], we specify fixed effects of *region* and *group*. We also specify two random effects of *participant_id* and *region.* This syntax specifies *1|participant_id* to model both overall individual differences as well as *1|region in participant_id* to model region-specific individual differences. Then, the emmeans package in R is used to extract marginal means from the model. The results of the pairwise comparisons [FDR - CTR] are presented below. SE = standard error; df = degrees of freedom.

| region | estimate | SE | df | t.ratio | p.value | fdr.adj.p |
|---|---|---|---|---|---|---|
| Posterior caudate | -0.179 | 0.144 | 26 | -1.238 | 0.227 | 0.453 |
| Posterior putamen | -0.045 | 0.144 | 26 | -0.314 | 0.756 | 0.756 |

**Table S5.** SCR data are fit using a linear mixed effects model in *N* = 28 and *N* = 69 individuals (after discarding 5 SCR recordings from the fMRI only arm). As per the equation [*lme(scr ~stimtype, random =~ 1|subid/run, data=filtered_data, method="REML")],* we specified the fixed effects of stimulus type (CS+ or CS-) and group. We also specify two random effects of *run (1, 2, 3, 4)* and participant_*id*. This syntax specifies *1|participant_id* to model both overall individual differences as well as *1|run in* participant_*id* to model run-specific individual differences. Because we are interested in main effects of stimulus type and group, we used the anova.lme function to extract main effects, presented below. numDF = numerator degrees of freedom, denDF = denominator degrees of freedom.

| N | Effect | numDF | denDF | F-value | p-value |
|---|---|---|---|---|---|
| 28 | Intercept | 1 | 111 | 13.80 | 0.0003 |
| | Stimulus type | 1 | 111 | 6.404 | 0.0128 |
| | Group | 1 | 26 | 0.752 | 0.3936 |
| 69 | Intercept | 1 | 275 | 21.62 | < 0.0001 |
| | Stimulus type | 1 | 275 | 12.02 | 0.0006 |
| | Group | 1 | 67 | 0.8112 | 0.3710 |

**Table S6.** Explicit recall data are fit using a linear mixed effects model in *N* = 28 and *N* = 74 individuals. As per the equation [*lme(Response ~stimtype + group + day, random =~ 1|subid/run, data=filtered_data, method="REML")],* we specify the fixed effects of stimulus type (CS+ or CS-), group and day (1, 2). We also specify two random effects of *run (1, 2, 3, 4)* and *participantt_id*. This syntax specifies *1|participantt_id* to model both overall individual differences as well as *1|run in participantt_id* to model run-specific individual differences. Because we are interested in main effects of stimulus type and group, we used the anova.lme function to extract main effects, presented below. numDF = numerator degrees of freedom, denDF = denominator degrees of freedom.

| N | Effect | numDF | denDF | F-value | p-value |
|---|---|---|---|---|---|
| 28 | Intercept | 1 | 334 | 115.4 | < 0.0001 |
| | Stimulus type | 1 | 334 | 246.1 | < 0.0001 |

| | | | | | |
|---|---|---|---|---|---|
| | Group | 1 | 26 | 0.089 | 0.7671 |
| | Day | 1 | 334 | 0.263 | 0.6083 |
| 74 | Intercept | 1 | 782 | 240.1 | < 0.0001 |
| | Stimulus type | 1 | 782 | 578.4 | < 0.0001 |
| | Group | 1 | 72 | 0.104 | 0.7485 |
| | Day | 1 | 782 | 0.045 | 0.8321 |

**Table S7.** Reaction time data during recall are fit using a linear mixed effects model in *N* = 28 and *N* = 74 individuals. As per the equation [*lme(Response ~stimtype + group + day, random =~ 1|subid/run, data=filtered_data, method="REML")],* we specify the fixed effects of stimulus type (CS+ or CS-), group and day (1, 2). We also specify two random effects of *run (1, 2, 3, 4)* and *participant_id*. This syntax specifies *1|participant_id* to model both overall individual differences as well as *1|run in participant_id* to model run-specific individual differences. Because we are interested in main effects of stimulus type and group, we used the anova.lme function to extract main effects, presented below. numDF = numerator degrees of freedom, denDF = denominator degrees of freedom.

| N | Effect | numDF | denDF | F-value | p-value |
|---|---|---|---|---|---|
| 28 | Intercept | 1 | 334 | 29.24 | < 0.0001 |
| | Stimulus type | 1 | 334 | 0.207 | 0.6491 |
| | Group | 1 | 26 | 0.008 | 0.9315 |
| | Day | 1 | 334 | 4.489 | 0.0348 |
| 74 | Intercept | 1 | 782 | 29.47 | < 0.0001 |
| | Stimulus type | 1 | 782 | 1.452 | 0.2285 |
| | Group | 1 | 72 | 0.144 | 0.7056 |
| | Day | 1 | 782 | 20.72 | < 0.0001 |

**Table S8.** fMRI contrast data during acquisition are fit using a linear mixed effects model *N* = 74 individuals. As per the equation *[lme(beta ~region*group+run, random =~ 1|participant_id, data=filtered_data, method="ML")],* we specify the fixed effects of region, group and run (1, 2,3,4). We also specify the random effect of *participant_id*. This syntax specifies *1|participant_id* to model overall individual differences. Then, the emmeans package in R is used to extract marginal means from the model. The results of the pairwise comparisons [FDR - CTR] are presented below only for significant parcels surviving FDR correction. Exploratory regressions are performed between the significant parcels and $\%\Delta BP_{ND}$ to rule out fMRI group differences on dopamine release. SE = standard error; df = degrees of freedom.

| contrast | Region (acronym) | estimate | SE | df | t.ratio | fdr.adj.p | Correlation with $\%\Delta BP_{ND}$ |
|---|---|---|---|---|---|---|---|
| fdr - hc | Area V6A (V6A) | 0.074 | 0.015 | 72 | 5.071 | 0.0006 | No |
| fdr - hc | Seventh Visual Area (V7) | 0.075 | 0.018 | 72 | 4.244 | 0.0063 | No |
| fdr - hc | Posterior InferoTemporal complex (PIT) | 0.063 | 0.018 | 72 | 3.560 | 0.0374 | No |
| fdr - hc | Eighth Visual Area (V8) | 0.062 | 0.018 | 72 | 3.516 | 0.0374 | No |
| fdr - hc | Medial amygdala (mAMY) | 0.061 | 0.018 | 72 | 3.439 | 0.0381 | No |
| fdr - hc | Fourth visual area (V4) | 0.059 | 0.018 | 72 | 3.348 | 0.0424 | No |
| fdr - hc | Area Lateral IntraParietal dorsal (LIPd) | 0.058 | 0.018 | 72 | 3.271 | 0.0432 | No |
| fdr - hc | Area V3CD (V3CD) | 0.057 | 0.018 | 72 | 3.235 | 0.0432 | No |
| fdr - hc | Ventral IntraParietal Complex (VIP) | 0.057 | 0.018 | 72 | 3.210 | 0.0432 | No |

**Table S9.** Predictions of the chaotic dopamine hypothesis using the posterior caudate striatal ROI.

| Prediction | Variables | Equation | **Statistic** ($\%\Delta BP_{ND}$) |
|---|---|---|---|
| ↓release → ↓BOLD for positive prediction error | $\%\Delta BP_{ND}$<br>ΔBOLD [CS+ - CS-] | ΔBOLD ~ $\%\Delta BP_{ND}$+Group | **t(25) = -2.65, p = 0.014** |
| ↓release → ↓autonomic responding | $\%\Delta BP_{ND}$<br>Skin conductance<br>Δ CS [CS+ - CS-] | Autonomic response ~ $\%\Delta BP_{ND}$+Group +Stimulus | t(25) = -0.21, p = 0.84 |
| ↓release → ↓valuation of stimuli | $\%\Delta BP_{ND}$<br>Explicit recall<br>Δ CS [CS+ - CS-] | Likert response ~ $\%\Delta BP_{ND}$+Group +Day(1+2) | t(52) = 0.47, p = 0.64 |
| ↓release → ↑negative symptoms | $\%\Delta BP_{ND}$<br>TEPS total | TEPS ~ $\%\Delta BP_{ND}$+Group | t(25) = -0.076, p = 0.94 |
| ↓release → ↑negative symptoms | $\%\Delta BP_{ND}$<br>TEPS anticipatory | TEPS ~ $\%\Delta BP_{ND}$+Group | t(25) = -0.723, p = 0.476 |
| ↓release → ↑negative symptoms | $\%\Delta BP_{ND}$<br>TEP consummatory | TEPS ~ $\%\Delta BP_{ND}$+Group | t(25) = 0.709, p = 0.485 |
| **↓release → ↓positive symptoms** | **$\%\Delta BP_{ND}$<br>paranoia checklist total** | **log(paranoia checklist) ~ $\%\Delta BP_{ND}$+Group** | **t(25) = 2.07, p = 0.049** |
| **↓release → ↓positive** | **$\%\Delta BP_{ND}$** | **log(paranoia checklist) ~** | **t(25) = 2.20** |

| symptoms | paranoia checklist frequency | %Δ$BP_{ND}$+Group | p = 0.037 |
|---|---|---|---|
| ↓release → ↓positive symptoms | %Δ$BP_{ND}$ paranoia checklist conviction | log(paranoia checklist) ~ %Δ$BP_{ND}$+Group | t(25) = 1.498, p = 0.147 |
| ↓release → ↓positive symptoms | %Δ$BP_{ND}$ paranoia checklist distress | log(paranoia checklist) ~ %Δ$BP_{ND}$+Group | t(25) = 1.977, p = 0.059 |

[9] Y. Zhang, J. Brady, S. M. Smith, and Fmrib, “An HMRF-EM algorithm for partial volume segmentation of brain MRI FMRIB technical report TR 01 YZ 1,” 2001.
